# Human gut microbiome gene co-expression network reveals a loss in taxonomic and functional diversity in Parkinson’s disease

**DOI:** 10.1101/2024.12.18.629142

**Authors:** Polina V. Novikova, Rémy Villette, Cédric C. Laczny, Brit Mollenhauer, Patrick May, Paul Wilmes

**Affiliations:** Luxembourg Centre for Systems Biomedicine, University of Luxembourg, Esch-sur-Alzette, Luxembourg; Department of Neurology, University Medical Center Göttingen, Göttingen, Germany; Paracelsus-Elena-Klinik, Kassel, Germany; Department of Life Sciences and Medicine, Faculty of Science, Technology and Medicine, University of Luxembourg, Belvaux, Luxembourg

## Abstract

Alterations of gut microbiome structure have been observed in a panoply of human diseases including neurodegeneration. However, the ecological and functional deficits of microbiome dysbiosis have yet to be understood. Here, using integrated multi-omics (metagenomics and metatranscriptomics), we resolve microbiome gene co-expression networks in individuals with Parkinson’s disease (PD) and healthy individuals. We uncover network modules with high closeness and degree centrality that represent core ecological functions, and identify key features lost in PD. More specifically, we observe a significant depletion in specific functions including secondary bile acid biosynthesis and flagellar assembly (FA) in PD. Strikingly, most hub genes, particularly those involved in bacterial microcompartments (BMCs) and FA, are predominantly found in healthy individuals. *Blautia* and *Anaerobutyricum* genera are the main contributors to these functions, showing significantly lower expression of BMC genes in PD. Additionally, we identify a strong correlation between the expression of BMC and FA genes, but also an apparent dysregulation in cross feeding between commensals in PD. Importantly, gene expression in PD was tied to reduced diversity in expressed genes, whereas in healthy individuals, higher expression levels were linked to higher diversity. Our findings reveal disruptions in key gut metabolic functions at both functional and taxonomic levels, potentially driving disease progression. Notably, we identify crucial microbiome-wide ecological features that should be restored in future gut microbiome rewilding efforts.

## Introduction

Parkinson’s disease (PD) is the second most prevalent neurodegenerative disorder, primarily caused by the loss of dopaminergic neurons and the formation of Lewy bodies in the brain as the major pathological hallmarks. PD is characterised by both motor and non-motor symptoms, with non-motor symptoms such as dysphagia, constipation, and bloating being linked to the gastrointestinal tract. Notably, idiopathic constipation, a common symptom of PD, often precedes motor symptoms by over a decade (Fasano et al., 2015), supporting the hypothesis that the disease can originate in the gut (Braak et al., 2003). Intestinal dysbiosis has been documented in PD (Cryan et al., 2020; Keshavarzian et al., 2015; Scheperjans et al., 2015), and evidence suggests that the gut might initiate or worsen the development of PD (Bhattarai et al., 2021; Hirayama & Ohno, 2021; Qian et al., 2018). Changes to gut microbiome structure can compromise gut permeability and the integrity of the intestinal barrier, affecting gastrointestinal epithelial cells, the immune system, and the enteric nervous system (Stolfi et al., 2022; Weiss & Hennet, 2017). Moreover, a dysbiotic gut is characterised by decreased microbial diversity (Weiss & Hennet, 2017). Multiple studies have uncovered evidence for dysbiosis in the gut microbiome of individuals with PD characterised by shifts in bacterial and archaeal taxa including *Akkermansia, Roseburia, Methanobrevibacter* (Boertien et al., 2019; Romano et al., 2021; Toh et al., 2022). In addition, we recently showed a decreased activity for *Roseburia sp.*, *Blautia obeum*/*wexlerae* and *Blautia massiliensis* in PD, a significant decrease in transcripts implicated in flagellar assembly and chemotaxis, and differences in the gut metabolome including decrease levels of chenodeoxycholic, glycocholic acid, glucuronic acid and glycerol, as well as an increase in β-glutamate, isovalerate and isobutyate (Villette et al., 2024).

Traditional bioinformatic and biostatistic methods frequently fall short of capturing the full complexity of microbiome dynamics and interactions. Indeed, differential expression analysis or multi-variate approaches do not capture the importance of functions within a complex ecosystem such as the gut microbiome. For that purpose, gene co-expression networks have been widely used to uncover functional gene clusters and pathways associated with various disease phenotypes (Cai et al., 2023; Meng & Mei, 2019). In this approach, co-expressed genes are classified into modules based on network properties, and highly connected genes are further identified as hub genes.

Here, we employ weighted gene co-expression network analysis (WGCNA) based on the ratio of metatranscriptomic (MT) to metagenomic reads transcript per million (MG TPM) from the gut microbiomes of individuals with PD and healthy individuals, referred here as healthy controls (HC). Using this approach, we managed to describe important shifts in the bacterial ecosystem metabolism, especially regarding bacterial microcompartments (BMCs), polyols transporters. Finally, we highlight the cross talk between commensal species and the decrease of taxonomic diversity of gene expression (tDGE) in PD compared to HC.

## Materials and Methods

### Patient, sample preparation, sequencing and data generation

All methods regarding patient recruitment, sample preparation, sequencing and data generation have been extensively described in a previous work (Villette et al, preprint link). Rapidly, the cohort included 29 healthy controls (HC), and 46 PD recruited at the Paracelsus-Elena Klinik, Kassel, Germany. An additional 20 HC were recruited as neurologically healthy subjects living in the same household as the PD participants. All subjects from both cohorts provided informed written consent, and the sample analysis was approved by the Comité National d’Ethique de Recherche of Luxembourg (reference no.: 140174_ND).

### Sample preparation

Fecal samples were collected at the clinics via a stool specimen collector (MedAuxil) and collection tubes (Sarstedt), as previously described (Heintz-Buschart et al., 2018). Samples were immediately flash-frozen on dry ice after collection. Samples were subsequently stored at –80 °C and shipped on dry ice. Extractions from fecal samples were performed according to a previously published protocol (Roume et al., 2012) conducted on a customized robotic system (Tecan Freedom EVO 200). Libraries were sequenced on an Illumina NextSeq500 instrument with 2×150 bp read length.

### Metagenomics and metatranscriptomics

For all samples, MG and MT sequencing data were processed and hybrid-assembled using the Integrated Meta-omic Pipeline (IMP) (Narayanasamy et al., 2016). Data was quality trimmed, adapter sequences were removed, MT rRNA reads were removed by mapping against SILVA 138.1 (RRID:SCR_006423) (Quast et al., 2013) and human reads were removed from MT and MG after mapping against the human genome (hg38) and transcriptome (RefSeq 212, RRID:SCR_003496). Pre-processed MG and MT reads were co-assembled using the IMP-based iterative hybrid-assembly pipeline using MEGAHIT (1.0.3, RRID:SCR_018551) (D. Li et al., 2015). After assembly, the prediction and annotation of genomic features such as open-reading frames (ORFs) and non-coding genes was performed using a modified version of Prokka (RRID:SCR_014732) (Seemann, 2014) and followed by functional annotation of those using Mantis (RRID:SCR_021001) (Queirós et al., 2021). Genomic features were quantified on MG and MT level using featureCounts (RRID:SCR_012919) (Liao et al., 2014) from the final gff file. Taxonomic annotation of reads and contigs was performed using Kraken2 (RRID:SCR_005484) (Wood et al., 2019) with a GTDB release207 database (RRID:SCR_019136) (http://ftp.tue.mpg.de/ebio/projects/struo2/GTDB_release207/kraken2) and a 0.5 confidence threshold.

### Co-expression network construction

Prior to network construction MG and MT count were normalised using transcript per million (TPM). We created normalised gene expression by using a ratio of MT TPM to MG TPM as described before (Roume et al., 2015). We then used the Python package WGCNA (RRID:SCR_003302) (PyWGCNA, version 2.0.4) to construct a co-expression network of KEGG orthologs (KO) normalised gene expression from the microbiome of individuals with PD and HC according to the WGCNA procedure (Rezaie et al., 2023). Normalised gene expression of KOs was power transformed prior to WGCNA, using PowerTransformer from *sklearn.preprocessing* (RRID:SCR_019053). The *WGCNA* function was run with the following parameters: minimum module size minModuleSize=20, dissimilarity threshold MEDissThres=0.18, networkType=’signed’. Modules were identified using average linkage hierarchical clustering and the dynamic tree-cut function.

### Topology metrics calculation

To analyse the structure of each network module, we calculated several key topological metrics. Module size was defined by the number of genes (nodes) in each module. Intramodular connectivity was measured as the sum of edge weights in the module adjacency matrix, divided by two. Mean connectivity represented the average edge weight within the module. Module diversity was assessed using the Shannon index, calculated on the sum of MT/MG ration of genes in a module (see “Diversity measures” for more detail). Centrality and clustering metrics were calculated using various functions from NetworkX (nx) (https://networkx.org<u>, RRID:SCR_016864</u>) (Hagberg A. et al., 2004), python package for the construction and analysis of complex networks. Specifically, clustering coefficient was calculated with *nx.average_clustering*, closeness centrality with *nx.closeness_centrality*, degree centrality with *nx.degree_centrality*, eigenvector centrality with *nx.eigenvector_centrality_numpy*, and betweenness centrality with *nx.betweenness_centrality*.

### Hub genes and intramodular hub genes selection

Hub genes are defined as the top 5% most connected genes present in trait-associated modules. For this purpose we used the 95^th^ percentile of connectivity for all modules with a significant trait-association using the *quantile()* function from R base package (RRID:SCR_001905). We also defined intramodular hub genes (iHub genes) by selecting 10% of the most connected genes from trait-associated modules.

### Selection of taxa of interest

In order to support the potential links between BMC and FA gene expression, we selected taxa of interest based on literature evidence of either flagellin expression, BMC formation and significant expression of either FA or BMC genes. In that manner we selected the taxa belonging to the following genera: *Agathobacter, Anaerobutyricum, Blautia, Faecalibacterium, Flavonifractor, Eubacterium, Lachnospira, Roseburia, RUG115* and all genera from *Lachnospiraceae*, *Oscillospiraceae* and *Ruminoncoccaceae* comprising “CAG” in their genus name.

### Diversity measures

#### Module diversity

We defined module diversity by the richness and evenness of normalised gene expression within a given module, whereby we summed normalised expression for each gene and applied the Shannon index from the vegan R package to the summed normalised expression (2.6.6.1, RRID:SCR_011950) (Oksanen et al., 2016). Hence, the module diversity index is calculated as following: *mH*′ = − ∑ *gi* log_*b*_ *gi*, where *mH*′ is the module diversity index and *gi* is the normalised expression of gene *i*.

#### Taxonomic diversity of gene expression

We defined tDGE as the diversity of taxa expressing a given gene using the Shannon index. Hence, the tDGE index is calculated as following: *gH*′ = − ∑ *si* log_*b*_ *si*, where *gH*′ is the tDGE index for gene *g* and *si* is the normalised expression of species *i* expressing the gene *g*.

#### Functional redundancy

Functional redundancy was calculated using the R package SYNCSA (1.3.4) using the function *rao.diversity()* (Debastiani & Pillar, 2012), functional redundancy is thereby defined as the difference between species diversity and the functional diversity (measured by Rao quadratic entropy) as proposed previously (de Bello et al. 2007).

#### Diversity of gene expressed per taxa

We quantified the diversity of gene expressed per taxa by summing the normalised expression, for each sample, at the genus level for the selected taxa (see above), *PeH17* and *Faecousia*. For that purpose, we used the observed number of genes expressed, the inverse Simpson index (vegan package) and Shannon index.

#### Statistics and plotting

All statistics and plots were handled in R (4.4.1) using Rstudio (2024.4.2.764, RRID:SCR_000432). Wilcoxon tests, spearman correlations tests and FDR corrections were performed using *rstatix* package (0.7.2, RRID:SCR_021240). Gene set enrichment analysis was conducted using fgsea R package (1.30.0, RRID:SCR_020938). Differential expression analysis was performed using DeSeq2 R package (1.42.1, RRID:SCR_015687). All plots were produced using ggplot2 (3.5.1, RRID:SCR_014601).

## Results

### Microbial co-expression network is linked to disease status

We constructed a network representation of the gut microbiome including samples from both HC (n=49) and PD individuals (n=46) using WGCNA (Langfelder and Horvath, 2008). For the multi-omics co-expression analysis, we used abundance-normalised gene expression data, more specifically the ratio of MT and MG transcripts per million (TPM, Figure 1) (Roume et al., 2015). From an original set of 11,876 microbial KO, referred in this paper as genes, we constructed a signed co-expression network of 4,789 genes. The co-expression network revealed 17 modules from which trait-association was found, four significantly associated with HC (M13, M2, M11 and M17, p < 0.05), five significantly associated with PD (M3, M6, M7, M8 and M15, p ≤ 0.05), and eight neither significantly associated with HC nor with PD (M1, M4, M5, M9, M10, M12, M14 and M16, Figure 2A).

**Figure 1.**
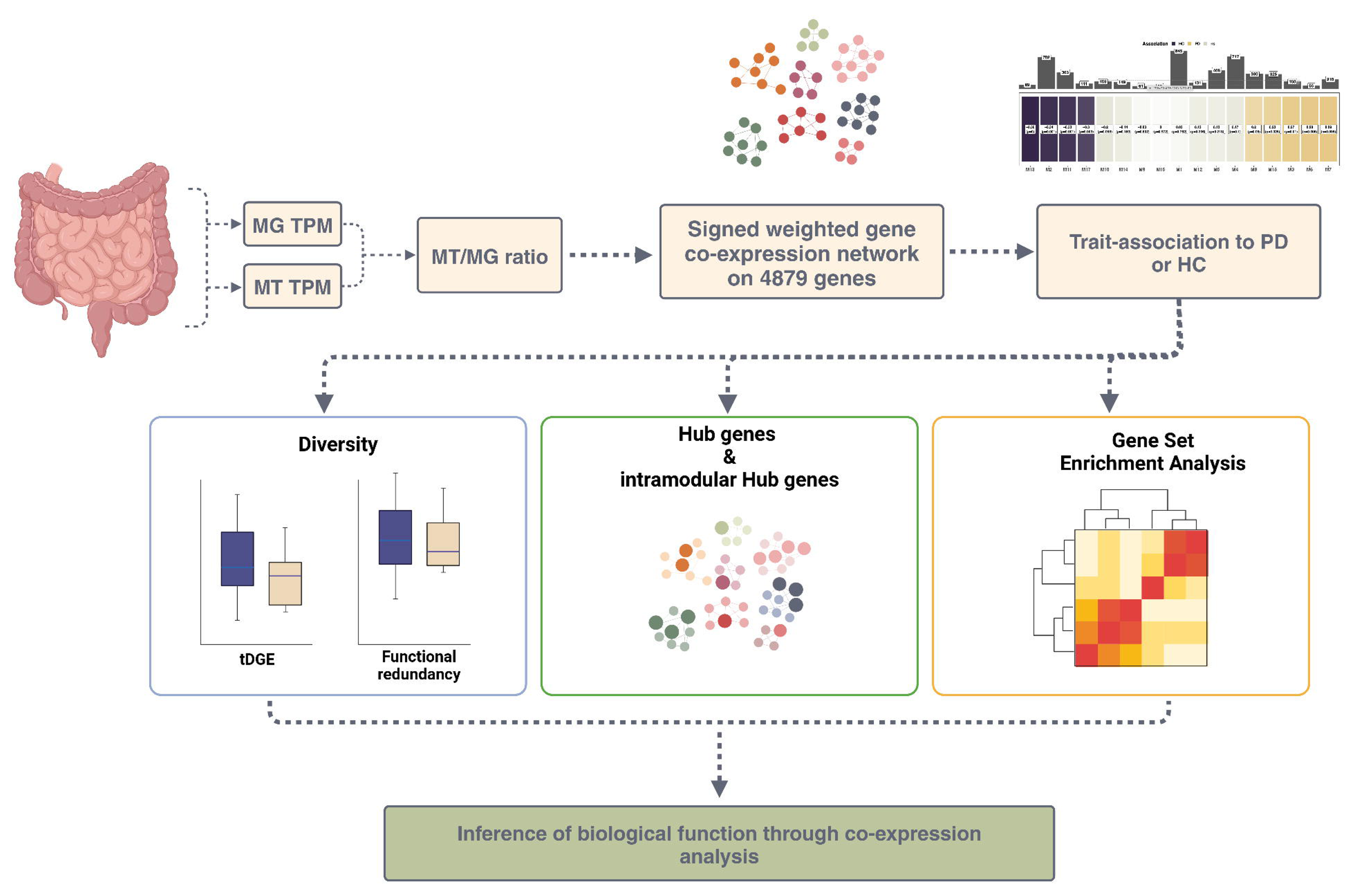
Overview of the data analysis workflow for identifying and analysing co-expressed gene modules using Weighted Correlation Gene Network Analysis. Metagenomic (MG) and metatranscriptomic (MT) counts per gene were converted into values representing the normalized gene expression MT/MG ratio. A co-expression gene network was then constructed based on a dataset of 4879 genes derived from both PD and HC individuals. This network revealed 17 distinct gene modules. Among these, modules significantly associated with either PD or HC were selected. Further analyses focused on gene diversity analysis, hub genes and gene set enrichments, aiming to uncover the ecological relevance of these modules in relation to the disease. Gene set enrichment analysis (GSEA), Parkinson’s disease (PD), healthy control (HC).

**Figure 2.**
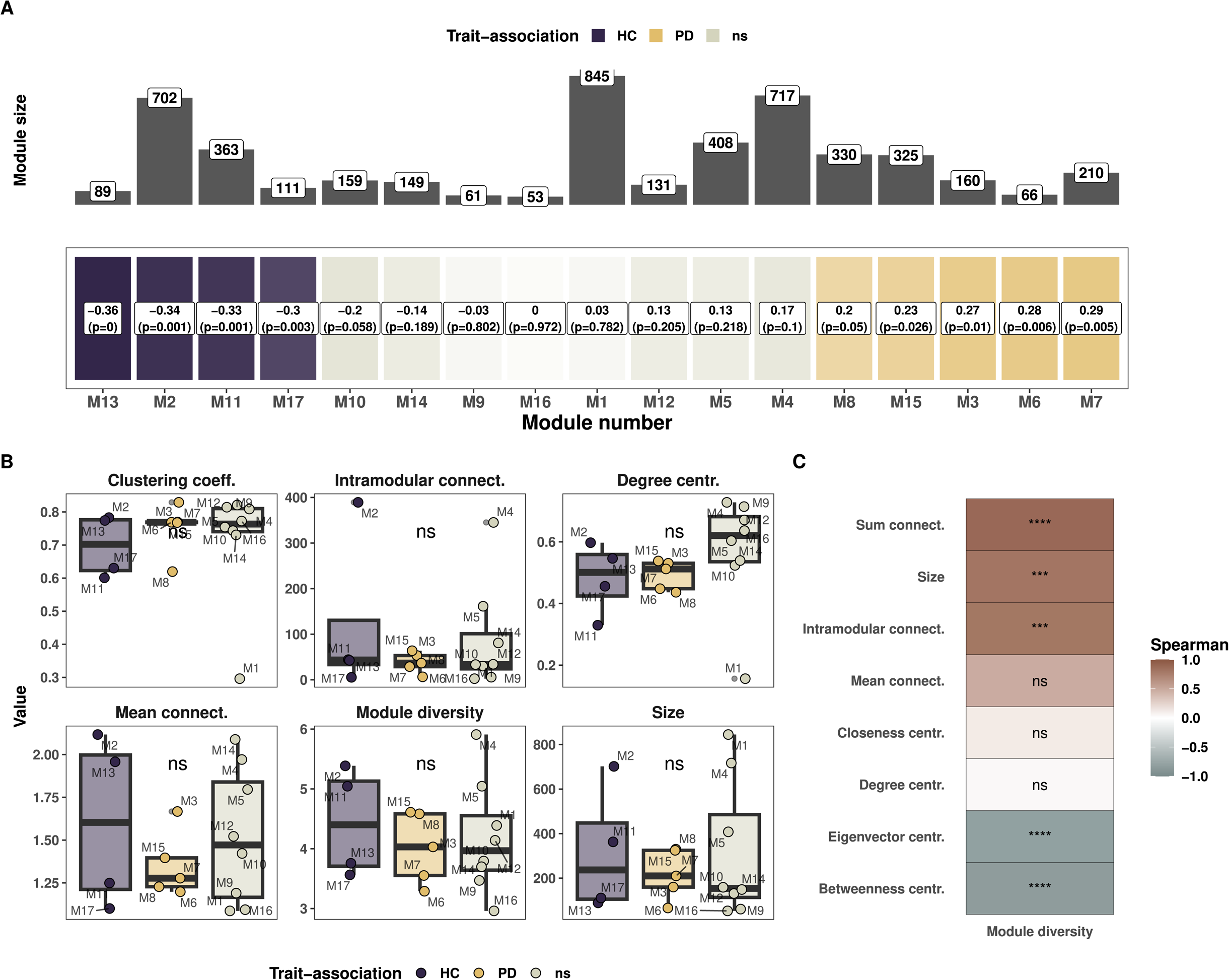
Weighted Correlation Gene Network Analysis reveals module associations with disease. **A.** Module trait relationship heatmap with correlation and p-values for each module. Modules are sorted from left to right (healthy to Parkinson’s disease) based on the eigenvector value. The top panel represents the number of genes belonging to each module. **B.** Network topology analysis for the modules grouped by trait association. A Kruskal and Wallis test was performed according to the trait association to compare modules based on their associations or not to one of the two groups. **C.** Correlation between module diversity and other topology features.

We next looked at topological features of the network and calculated the module diversity by defining the diversity of genes found within each module, using the Shannon index (Shannon, 1948). Modules M2 and M4 showed high mean connectivity, highest intramodular connectivity and highest module diversity despite their large size (Figure 2B and Supplemental fig. 1). M1 had the lowest clustering coefficient, degree centrality, eigenvector centrality and closeness centrality, thereby demonstrating that M1 acted as a default module, comprised of genes with the lowest clustering coefficient in the dataset (Figure 2B and Supplemental fig. 1). M6 exhibited high values of betweenness and eigenvector centrality but low connectivity (Figure 2B and Supplemental fig. 1A). When analysing the modules based on trait association, we found no statistically significant differences in topological measures but important variation between the modules, irrespective of trait association (Kruskal and Wallis test, p > 0.05, Figure 2B and Supplemental fig. 1A). However, M1 being a clear outlier for clustering coefficient, degree and closeness centrality, we removed it to re-assess modules based on trait association and did indeed find a significantly higher degree and closeness centrality (Kruskal and Wallis, p = 0.023 and p = 0.013, data not shown). Finally, it was apparent that module diversity and the different topology metrics such as the sum of connectivity and intramodular connectivity were correlated (R^2^ > 0.8, p < 0.001, Figure 2C). In contrast, eigenvector and betweenness centrality were anticorrelated with module diversity (R < −0.8, p < 0.001, Figure 2C). Overall, we found significant anti-correlations between connectivity measures and closeness/betweenness centralities while the clustering coefficient was correlated to degree/eigenvector centrality (Supplemental fig. 1C).

### Modules associated with healthy individuals exhibit enrichment in flagellar assembly and secondary bile acid biosynthesis

We next proceeded to gene set enrichment analysis (GSEA) to resolve the pathway-level information on the modules. Based on KEGG pathways, we found that on average 43.7% (min: 26%, max: 59%, Figure 3A) of genes within modules did not belong to any pathway. With respect to known pathways, modules were comprised on average of 28.5 different pathways ranging from 10 (M6) to 42 pathways (M1) (Supplemental fig. 2). We identified an enrichment in flagellar assembly in M13 and secondary bile acid biosynthesis in M11, which, in both cases, were associated with HC. We also found an enrichment in genes involved in biofilm formation for M16 (no trait association) and no statistically significant enrichment in PD-associated modules (Figure 3, q < 0.05, q < 0.05 and q > 0.05, respectively, GSEA). Although not statistically significant after correction, we noticed enrichments in the following pathways within modules associated with PD: glycerolipid metabolism (M3), peptidoglycan biosynthesis (M15), lipoic acid metabolism and valine degradation (M7) (Figure 3B, p < 0.01). We also noticed the presence of beta-lactam resistance genes (*oppA, oppB, oppC, oppD* and *oppF*) and quorum sensing genes in PD-associated module M6 (Supplemental fig. 1). In addition, genes involved in methane metabolism were present in the PD-associated modules M6 and M8 (Supplemental fig. 1).

**Figure 3.**
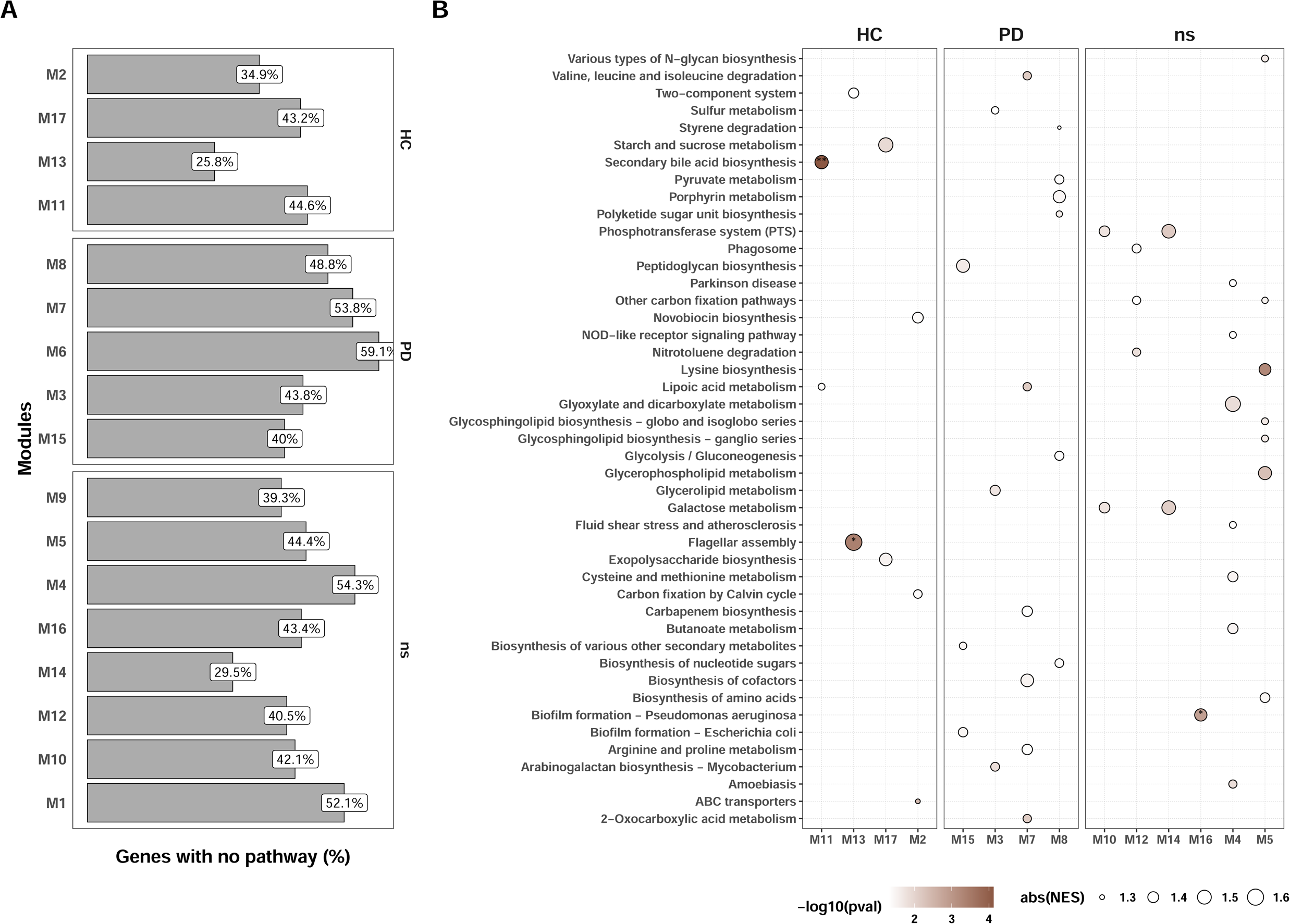
Gene set enrichment analysis on KEGG pathway highlights enrichment in HC modules but not in PD modules. **A.** Count of genes within a module with undescribed pathways or not belonging to any KEGG pathway. **B.** Gene Set Enrichment Analysis. Dots reflect significant enrichment (p < 0.05), colored by −log10(p-value). Asterisks represent significant enrichments after FDR correction (q < 0.05).

### Hub genes are restricted to health-associated modules

We next defined hub genes to assess key functions within the microbial co-expression network. We first selected the 95^th^ percentile of genes to select the top 5% of genes with the highest connectivity from all modules associated with HC and PD, excluding genes from modules not associated with either of the groups. Out of the 126 genes, 120 were from HC-associated modules and 6 from PD-associated modules (Figure 4A). To resolve more genes from PD-associated modules, we selected 10% of the most connected genes within each module, representing here intramodular hub genes (iHub genes), thereby retrieving 125 from HC-associated modules and 108 from PD-associated modules (Figure 4A). For the 95^th^ percentile approach, most genes belonged to module M2 (82%) and the rest belonged to M13 (13.5%), M3 (3.9%) and M15 (0.8%, Figure 4B). Hub genes associated with HC were mainly involved in energy production (oxidative phosphorylation, glycolysis/gluconeogenesis), transporter activity (ABC transporters), nucleotide metabolism (pyrimidine and purine metabolism), saccharide metabolism (pentose and glucuronate interconversions, starch and sucrose metabolism) as well as microbial motility (flagellar assembly and two-component system) (Figure 4C). With the 10% per module approach, we noted the presence of two glutamate synthases (*GLT1* and *gltB*) in the module M11 linked to alanine, aspartate and glutamate metabolism (Figure 4C).

**Figure 4.**
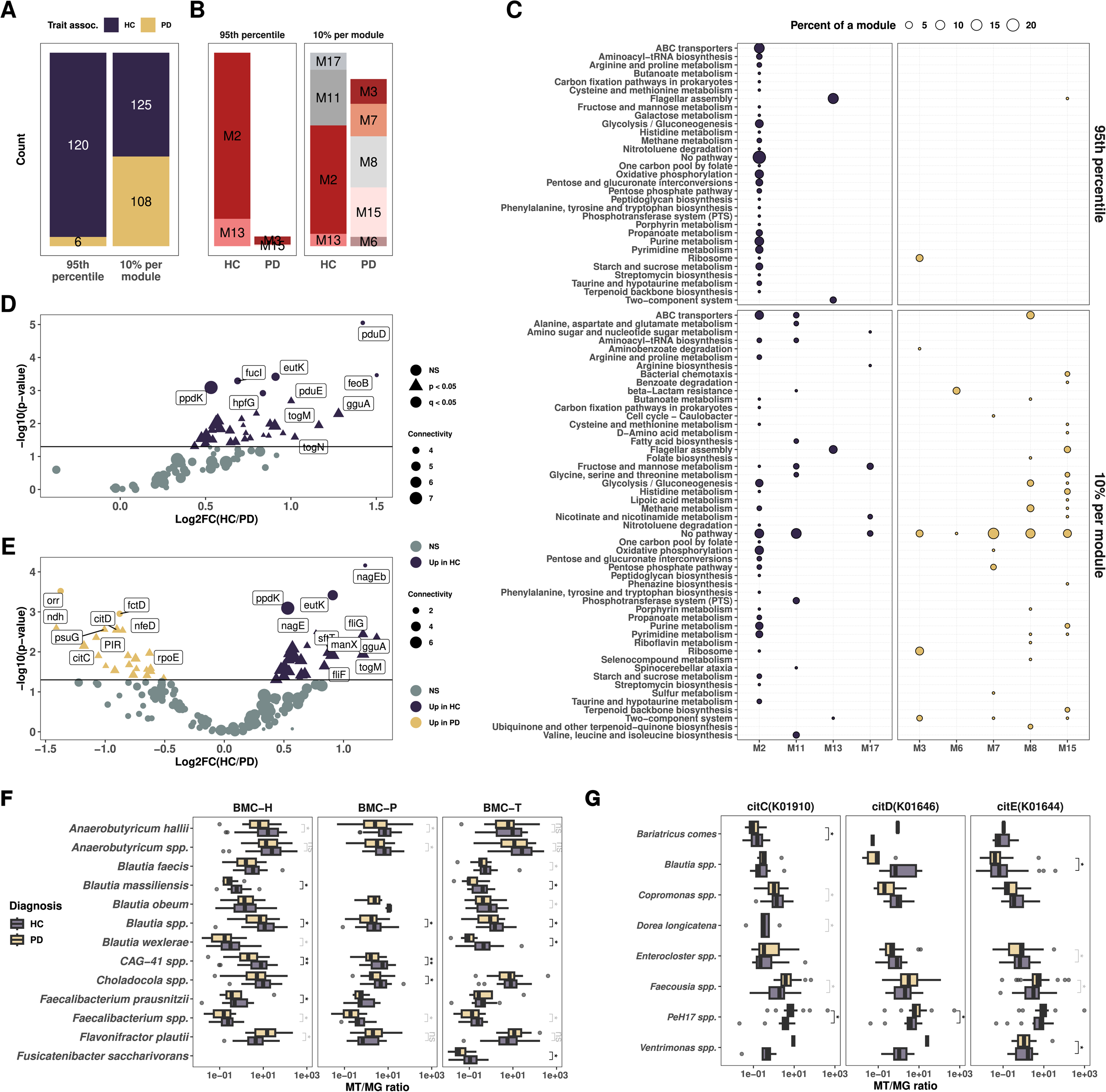
Hub genes are mostly associated with healthy individuals. **A.** Bar plot highlighting the counts of hub genes selected based on top 100 connected genes (left panel) and 10 % top connected genes per module (right panel). **B.** Dot plot representing the count of pathways per module for the hub genes and iHub genes. The size of the dots represents the proportion of a given pathway within a module. **D and E.** Volcano plots of differential expression of genes for the hub genes selected with the 95^th^ percentile connected genes in the network (**D.**) and top 10% connected per modules (**E**). Dots are colorized by group and shaped on the level of significance, triangular shape for p < 0.05 and round colored shape for q < 0.05. **F and G.** Boxplots representing gene normalized expression resolved at the species level for the BMC shell proteins (**F**.) and citrate lyases (**G**.). All tests are Mann-Whitney tests with q values (FDR corrected) depicted in black and p values (q > 0.05 after FDR correction) depicted in grey.

Amongst the hub genes, we found a significant increase in their expression in HC including flagellar assembly (*flgB, fliQ, fliS, flgE, fliK, flgL, fliD, fliP,* p < 0.05, Figure 4D). We noted that genes involved in bacterial BMC catabolism and metabolism (eut and pdu operons) were significantly increased in HC (*eutL, pduE, eutK,* q < 0.05, DeSeq2, Figure 4D and E) along with genes involved in polyols transport (togB, togM, togN, gguA, lacF and ganA, p < 0.05, q > 0.05, DeSeq2 Figures 4D and E). Of note, the togBMNA operon is an important transporter for oligogalacturonide, a by-product of pectin degradation and involved in short chain fatty acid (SCFA) production. We also noted a significantly increased expression for citrate lyase genes in individuals with PD (*citD, citC* and *citE,* p < 0.05, q > 0.05, Figure 4E).

We then proceed to taxonomically resolve the expression of these genes. First, we noted the variability in ortholog annotations between the pdu and eut operons, these annotations were dependent on the taxonomy. Indeed, most of the shell proteins annotated as members of the eut operon, while catabolism/anabolism genes were annotated as members of the pdu operon. We manually grouped the genes according to their described functions in the literature. Resolving the gene expression at the species level revealed a significant decrease in expression for these genes in PD, including in *Blautia* spp.*, Blautia massiliensis, Blautia obeum*, *Blautia massiliensis, CAG-41 spp., Anaerobutyricum soehngenii* and *Faecalibacterium prausnitzii* (q < 0.05, Mann-Whitney test, Figure 4F and Supplemental fig. 3A). Of note, this decrease was also observed at the genus level (data not shown). Interestingly, we found an increase in *Flavonifractor plautii*’s expression of BMC genes encoding for BMC-H and pduQ in PD (p < 0.05, Mann-Whitney test, Supplemental fig. 3A). In addition, we noted a decrease of expression in for togBMN genes in *F. prausnitzii* and *Gemmiger qucibialis,* for lacFG, gguA and wzm in *Blautia spp.*, and for ganPQ and msmX in *Agathobacter spp.* (q < 0.05, Mann-Whitney test, Supplemental fig. 3B-C). Finally, citrate lyase genes expression revealed the increased expression of *PeH17 spp.* and *Faecousia spp*. in individuals with PD (q < 0.05 and p < 0.02, respectively, Wilcoxon test, Figure 4G).

### Bacterial microcompartment genes are correlated with flagellar expression

We next investigated the links between the expression of BMC genes and FA genes. For this, we tested the correlation between these genes using both normalised gene expression (reminder: MT/MG ratio) and MT TPM. In addition, we tested the correlations for all the genes from all the taxa or selected taxa based on literature evidence and evidence in this work (see material and methods for details). We found strong correlations between the levels of gene expression for BMC and FA genes (Figure 5A and Supplemental fig. 4A-B). Indeed, when using all the taxa from the dataset we noted 749 significant correlations before FDR correction and 139 significant correlations after FDR correction when using normalised gene expression (Figure 5A, p < 0.05, q < 0.05, respectively, Spearman correlation test); 831 significant correlations before FDR and 289 significant correlations after FDR correction when using MT TPM (Figure 5A, p < 0.05, q < 0.05, respectively). In addition, using only the selected taxa resulted in even more significant correlations both for normalized gene expression (444 genes with p < 0.05, 338 genes with q < 0.05, respectively, Spearman correlation test) and MT TPM (806 genes with p < 0.05, 777 genes q < 0.05, respectively, Figure 5A). Next, we specifically checked the correlation of hub genes for the selected taxa, and we found significant correlations between hub genes from BMCs and FA genes both for normalized expression and MT TPM (Figure 5B and C, q < 0.05, Spearman correlation test). Finally, we investigated the shared presence of the different processes in the identified taxa. Interestingly, we found that *Blautia*, *Anaerobutyricum* and *Flavonifractor* were expressing mainly ABC transporters and BMC genes while *Roseburia*, *Lachnospira* and *Eubacterium* expressed almost no BMC genes (Figure 5D). We also noted that many cryptic genera, such as CAG-115, did not express BMC genes (Figure 5D). The correlations between BMC and FA genes are highlighting a cross-feeding process between the commensals.

**Figure 5.**
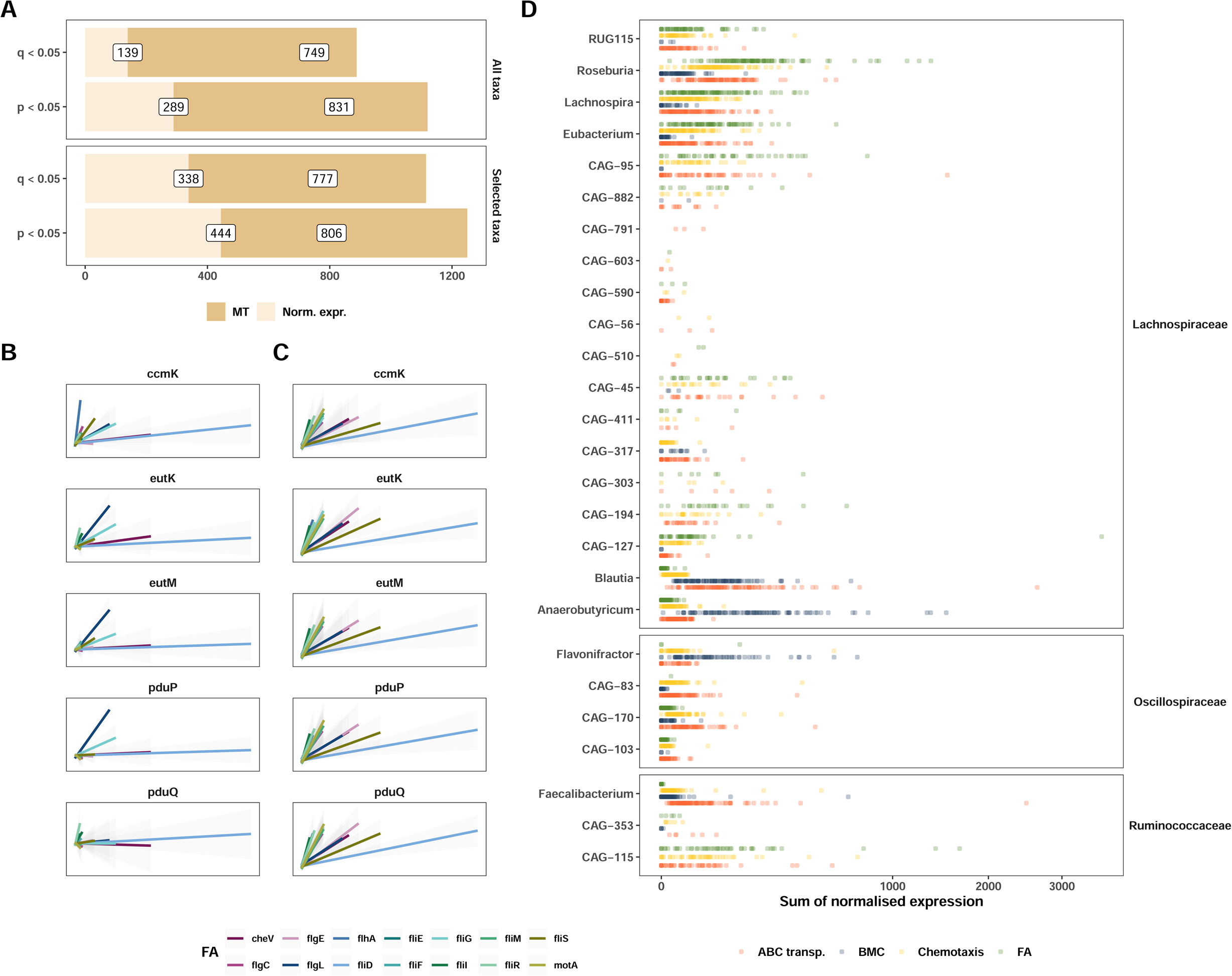
Bacterial microcompartments are correlated with genes involved in flagellar assembly. **A.** Bar plots highlighting the number of positive correlations before and after FDR correction for normalized expression and MT TPM when taking all taxa expressing the BMCs genes and FA genes (upper panel) and relevant taxa from figure 4 (lower panel). All tests are based on the Spearman correlation. **B.** Correlation plots including selected taxa, both for normalized expression and MT TPM, considering only hub genes from the 10% per modules approach. Tests are based on the Spearman correlation and all correlations are significant after FDR correction (q < 0.05). **C**. Dot plot representing the sum of normalised expression per sample for a given process. BMC, FA and Chemotaxis gene grouping comprise all genes detected in the dataset, while ABC transporters only include the following genes: togBMNA, lacDEFGAICR, man, gguAB, ganAQ, msmEFGX, and wzm. Genera represented here are the “selected taxa” presented in A and B.

### Genes enriched in Parkinson’s disease exhibit lower taxonomic diversity of gene expression

After noting interesting modifications in the co-expression network and the lack of hub genes in PD-associated modules, we decided to investigate functional redundancy and the taxonomic diversity of gene expression (tDGE) to assess the impact of taxonomic differences at the level of gene expression. Here we define tDGE as the diversity of taxa expressing a specific gene, while functional redundancy denotes the measure of taxonomic and functional diversity present within a sample (Tian et al., 2020). Our subsequent analyses aimed at differentiating between genes found in module association to either one of the conditions (trait-association, Figure 6A-C) and genes with differential expression showing an increase or decrease in PD (Figure 6D-F). We did not observe a difference in functional redundancy between HC and PD individuals (Figure 6A, p > 0.05, Wilcoxon test). Interestingly, we found no significant differences for overall tDGE for non-iHub genes but a significantly lower tDGE for iHub genes in PD (Figure 6B, Mann-Whitney test, p > 0.05 and p < 0.01, respectively). We also noticed iHub genes had a higher tDGE than non-iHub genes (Figure 6C, Wilcoxon test, p < 0.001). We next compared the differential expression and functional diversity of a given gene. We found that gene expression was linked to either an increase in tDGE (more microbes expressing a given gene) or a loss in tDGE (less microbes expressing a given gene) (Figures 6D and E).

**Figure 6.**
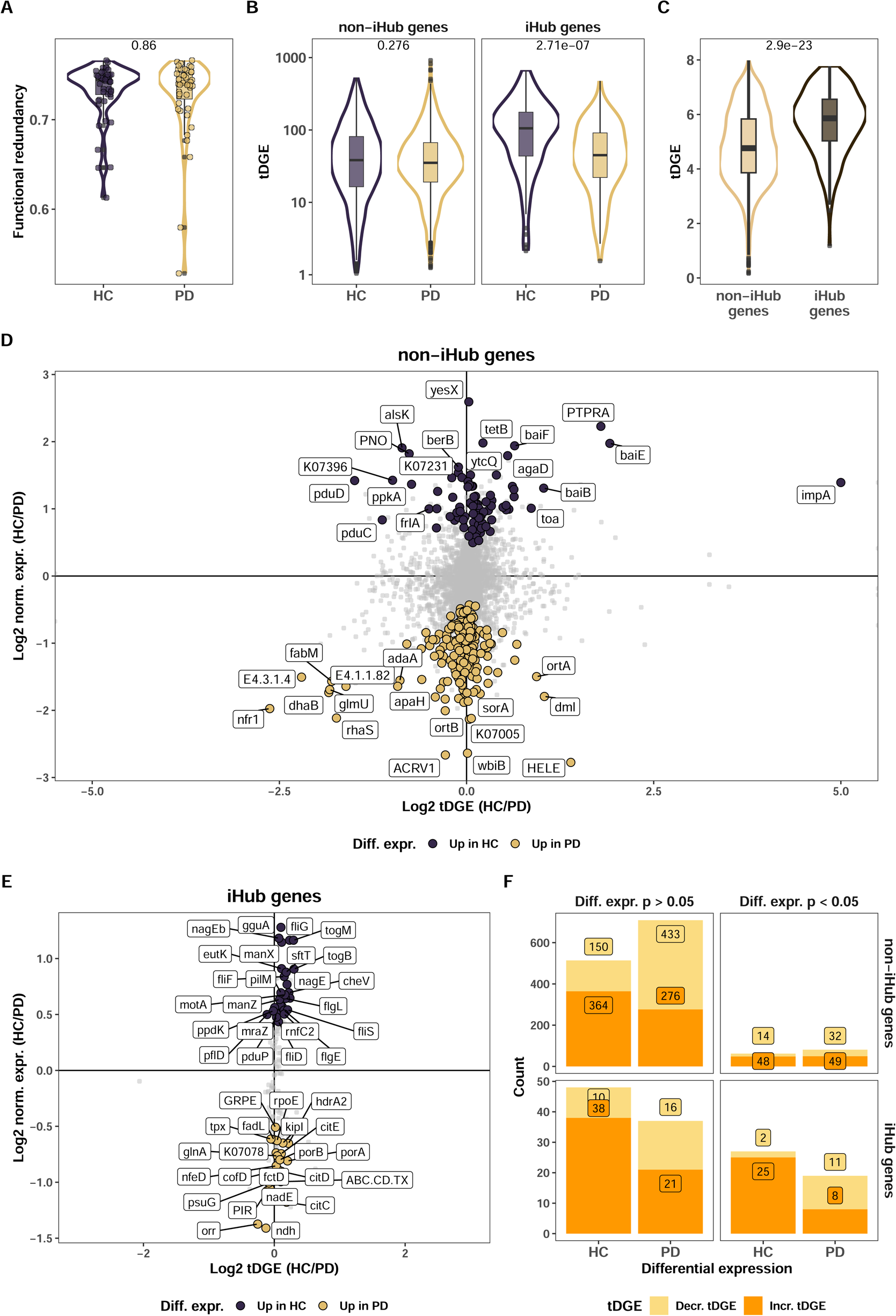
Gene diversity is decreased in PD. **A.** Boxplot representing functional redundancy for each sample according to disease status. **B.** Boxplot representing gene expression diversity according to disease status. **C.** Boxplot representing gene expression diversity grouped by hub genes belonging or not. **A**, **B** and **C** figures represent p-values from Mann-Whitney tests. **D and E.** Differential abundance versus gene expression diversity for a given gene for non-Hub genes (**D.**) and Hub genes (**E.**). Y-axis represents log2-fold change of normalized expression and X-axis the log 2-fold change of gene expression diversity. Dots are labelled and coloured for genes with p-value < 0.05. **F.** Stacked bar plot representing the counts of genes with an increased or decreased tDGE for iHub and non-iHub genes. Genes are classified into the PD or HC groups according to the sign of log2FC of normalized expression and faceted according to DEG significance.

We were able to categorize genes into two groups: those with increased expression linked to a higher tDGE, and those with increased expression linked to a lower tDGE. We finally looked at the proportion of genes within these two above-mentioned categories. We found that a significantly higher proportion of genes with decreases in tDGE in PD compared to HC for both hub genes and non-hub genes that had significantly different expression (p = 0.0004 and p = 0.049, Fisher exact test and Chi-square test, respectively, Figure 6F). Of note, we noted the same link between tDGE and differential expression in PD for genes that did not reach significance (Figure 6F). Interestingly, iHub genes with significantly increased differential expression in PD had a higher fraction of decreased tDGE than non-iHub genes, the difference was however not significant (57% and 43%, respectively, p=0.23, Chi-square test, Figure 6F).

Finally, we assessed the diversity of functions expressed in the selected taxa, *PeH17* and *Faecousia*. We found a consistent decrease in diversity for the genes expressed by *Roseburia* and *Eubacterium* genera by considering numbers of distinct genes expressed, the inverse Simpson index and the Shannon index (p <0.05, Mann-Whitney test, Supplemental fig. 5). Additionally, we also found a statistically significant decrease for *Lachnospira* and *Blautia* genera when quantifying the numbers of genes expressed (p < 0.05 and p < 0.01, Mann-Whitney test, respectively, Supplemental fig. 5). We also noted an increase in the inverse Simpson index for *Faecousia* and in the number of genes expressed by *PeH17*. However, these increases did not reach statistical significance (p = 0.084 and p= 0.082, respectively, Mann-Whitney test, Supplemental fig. 5).

## Discussion

Most knowledge on microbiome perturbations and associations with diseases comes from either 16S rRNA gene or whole genome sequencing which do not allow the ability to assess transcriptional activity of the microbiome. We previously demonstrated significant alterations of the gut microbiome in individuals with PD and highlighted strong differences in the expression of flagellar assembly and chemotaxis genes (Villette et al., 2024). Here, we aim at analysing the dataset using a system ecology approach to uncover microbial key regulator functions associated with PD. More specifically, we employ a co-expression network-based approach on MT and MG data to assess the microbiome-wide impacts of these changes. We identify strong associations between co-expression network modules and disease status, with four modules linked to HC and five to PD. Additionally, we find eight modules that were not associated with either HC or PD. Except for module M1, that is mainly a module comprised of genes not clustering with other genes, we find higher values of degree and closeness centrality for non-associated modules compared to the trait-associated modules. Given their overall high centrality and the lack of trait association, these modules may represent stable, core functions that support the integrity of microbial ecosystem services in both health and disease.

Hub genes represent biologically relevant properties and are more likely to reflect key functions that might be lost in diseased individuals such as PD (Calabrese et al., 2012; Horvath et al., 2006; Langfelder et al., 2013; Zhang & Horvath, 2005). Therefore, identifying hub genes is a valuable strategy to describe key functions that define healthy versus diseased individuals. Using this approach, we first select the 95th percentile of connected genes from modules associated with either HC or PD to uncover the most important genes in the co-expression network. This strategy reveals that 95% of the hub genes are associated with HC. To gain more insight into the PD-associated microbiome, we select the top 10% most connected genes per module, defined here as iHub genes. With this approach, we uncover citrate lyase as being enriched in PD. We are able to link the taxonomic expression of these genes to *PeH17* spp. and *Faecousia* spp. Interestingly, *PeH17* along with *Christensenella* are part of the *Christensenellales* order, the latter is usually described as increased in PD (Cirstea et al., 2020; Huang et al., 2023) and cross-feeding with *Methanobrevibacter* (Ruaud et al., 2020). Of note, *PeH17* is poorly characterised, and we currently do not know if *PeH17* exhibits the same cross-feeding activities with *Methanobrevibacter*. To our knowledge, there is no record of citrate lyase genes being associated with disease in humans and so far, no evidence of *PeH17* being associated with a disease in humans. Interestingly, thirteen genes from the flagellar assembly pathway are iHub genes and nine when using the 95^th^ percentile approach, showing once more the importance of this pathway in the microbial network, especially in the context of PD. Finally, the high number of hub genes from HC-associated modules is noteworthy, as it indicates that there is clear shift in the expression of key functions in the gut microbiome of PD.

Amongst modules, we find that M2 (HC-associated module) is of particular interest based on its connectivity, module diversity and centrality. M2 comprises the most connected genes and especially genes from the pdu and eut operons, two operons involved in the assembly of bacterial microcompartments. These operons are responsible for the utilization of ethanolamine and 1,2-propanediol, an important energy source for bacteria and typically associated with pathogenic bacteria (Ravcheev & Thiele, 2014; Tsoy et al., 2009), as they confer a growth advantage by utilizing abundantly present 1,2-propanediol and ethanolamine (EA) (Dank et al., 2021; Vance, 2018). However, it has been recently described that a wide range of commensals are also expressing these genes (Asija et al., 2021; Jallet et al., 2024; Q. Li et al., 2024; Reichardt et al., 2014). We find that the expression of these genes is decreased in PD compared to HC, especially in genera such as *Blautia* and *Anaerobutyricum* but increased in *Flavonifractor plautii*, a bacterial species that we previously highlighted as associated with HC (Villette et al., 2024, preprint link). In addition to BMC genes, we find genes from the togBMNA operon as hub genes and with genes significantly decreased in PD. These genes are part of ABC transporters involved in pectin degradation and production of short chain fatty acids (SCFAs) (Yüksel et al., 2024). We also note the presence of lacFG, ganPQ, wzm/wzt and other members of ABC transporters responsible of polysaccharides/polyols transport. Resolving the corresponding taxa reveals that these genes are significantly downregulated in members of *Faecalibacterium*, *Gemmiger*, *Roseburia* and *Blautia* genera. Along with the decrease in expression of BMC genes in PD, important metabolic processes, such as polyols transport (galacturonides, rhamnose, lactulose, etc.), are clearly downregulated in PD gut microbiome. Gut commensals lose keystone functions in the context of PD, resulting in the apparent loss of a gut microbial functional equilibrium.

In addition to the functional shifts seen in PD, we highlight strong correlations between the expression of BMC and FA genes even though these genes are not expressed by the same taxa. Indeed, BMC genes are mainly expressed by *Blautia, Anaerobutyricum* and *Faecalibacterium* species while FA are mainly expressed by *Roseburia, Lachnospira* and *Eubacterium* species. Interestingly, a recent study reported that GH26 b-1,4-endomannanase, a glycosyl hydrolase present on the surface of *R. intestinalis* but absent in *F. prausnitzii*, was necessary to grow on gluco-galactomannans (Lindstad et al., 2021). Our observation of reduced expression levels of BMC genes and polyol transporters in genera such as *Blautia*, *Anaerobutyricum*, and *Faecalibacterium*, along with the downregulation of FA, suggests a decline in cross-feeding interactions amongst these commensals in the gut microbiome of PD. However, it remains unclear whether the decreased expression of FA precedes or follows the reduced expression of BMC genes and polyol transporters. Interestingly, *Faecalibacterium*, *Blautia* and *Roseburia* species abundances are also decreased in other diseases such as IBD (Machiels et al., 2014), cancer (Wirbel et al., 2019; Ye et al., 2023; Yi et al., 2021; Yu et al., 2024) or Type 2 diabetes (Guo et al., 2024), underpinning the importance of these specific taxa.

In addition to the functional shifts, we observe a decreased tDGE in PD gut microbiome but no differences in overall functional redundancy. However, and crucially, we show here that most of the genes overexpressed in PD are linked to a decrease in the diversity of taxa expressing these functions. Interestingly, this decrease was even more apparent for genes that we defined as iHub genes, highlighting the reduced expression of keystone genes, such as BMC genes, in PD linked to keystone taxa of the gut microbiome, most notably *Blautia* species. In contrast to PD, in HC-associated genes, we note an opposite relationship, whereby an increased gene expression is linked to increased tDGE. Moreover, we found a decrease in functionality at the genus level. Indeed, *Roseburia*, *Blautia* and *Eubacterium* are expressing fewer genes in PD. On the other hand, we show here that *Faecousia* and *PeH17*, even if not reaching statistical significance, exhibit an increased diversity in genes expressed in the PD gut microbiome. All in all, this is to our knowledge, the first account of a decreased diversity in transcriptomic expression and functionality in the PD gut microbiome, bringing a new functional and ecological understanding to the concept of human gut microbiome dysbiosis.

Using the present study design, we were unable to measure resilience and stability of the gut microbiome in PD in comparison to healthy individuals as these measures are time-dependent. Nonetheless, a general decrease in taxa expressing key functions should result in, or be secondary to, decreased resilience and stability, as previously hypothesised (Ives & Carpenter, 2007; Loreau & Behera, 1999). Future studies should aim at characterising the gut microbiome in a longitudinal manner to evaluate the dynamics of the gut microbiome in a disease context. It is important to note that PD is a late onset and long-lasting progressive disease, understanding early perturbations in the gut ecosystem would enable assessment of the association between the gut microbiome and PD progression.

So far, little is known about the dynamics of the gut microbiome and the possibility of recovering from a dysbiotic gut microbiome in PD and during PD progression. In the same line, the dynamics and resilience of faecal transplantations, an emerging therapeutical option to recover from microbiome dysbiosis, are poorly understood. Human to mouse or human to organ-on-chip faecal transplantations studies are required to test the possibility to recover keystone functions from a dysbiotic gut microbiome.

In conclusion, we report here the importance of studying the functional capacity and diversity in the gut microbiome of individuals with PD. We see here, an overall decreased taxonomic diversity of gene expression and a decreased of functional diversity for keystone taxa. We also highlight de dysregulation of the cross-feeding of the gut microbiome in health and disease. *Faecalibacterium*, *Blautia* and *Roseburia* species are distinguishing themselves as keystone species in the healthy gut microbiome. Efforts are required to deepen our understanding of the interdependencies among these species, as well as to investigate ways to stimulate the functions of these taxa. Ultimately, in alignment with the ecological principles, researchers and clinicians should focus on rewilding the gut microbiome’s functions in diseased individuals to restore the diversity of functions within the gut microbiome ecosystem.

---

**Supplemental fig. 1. Additional topology metrics from WGCNA.**

**A.** Boxplots representing betweenness, closeness and eigenvector centrality. Kruskal and Wallis test. **B.** Correlation between topology metrics and module diversity. All tests are based on the Spearman correlation. **C.** Heatmap representing the correlation between various topology metrics. Tests are Spearman correlation test.

**Supplemental fig. 2. KEGG pathways per module.**

Counts of KEGG pathways per module for the hub genes. The size of the dots represents the proportion of a given pathway within a module.

**Supplemental fig. 3. BMC metabolism and ABC transporter gene expression resolved by taxa.**

**A.** Normalised gene expression for BMC metabolism, genes are grouped based on the described enzymatic activities. **B.** Normalised expression for togBMN genes for bacterial species with significant differential expression. **C.** Normalised expression for additional ABC transporter genes found as hug genes, for bacterial species with significant differential expression. All tests are Mann-Whitney tests corrected with FDR.

**Supplemental fig. 4. BMC genes correlate with flagella assembly genes.**

Heatmaps representing spearman correlation coefficients between genes involved in BMC formation, catabolism or anabolism and genes involved in flagellar assembly. **A.** Heatmap correlation tests for selected bacteria using normalized expression. **B.** Heatmap correlation tests for selected bacteria using MT TPM.

**Supplemental fig. 5. Diversity of genes expressed by *Roseburia*, *Blautia* and *Eubacterium* genera is decreased in PD, but increased for *Faecousia*.**

Alpha diversity measures of gene expressed in each genus. Plot are representing the sum of normalised expression for each sample and for each selected genus (see Material and Methods), with the addition of the *PeH17* and *Faecousia* genera. Values are representing the different diversity measures: inverse Simpson index, observed number of genes and Shannon index. All p-values are from a Mann-Whitney test.

## Data availability

Due to privacy restrictions, the datasets generated by this study are available upon reasonable request by contacting Paul Wilmes.

## Code availability

The IMP pipeline, which was used for analysis of metagenomic and metatranscriptomic data, is available at https://gitlab.lcsb.uni.lu/IMP/imp3. The R and python code used for statistical analyses and visualizations is available at https://gitlab.lcsb.uni.lu/ESB/[TBA].

## Acknowledgments

We thank the staff of the Luxembourg Centre for Systems Biomedicine (LCSB), particularly the sequencing platform and the metabolomics platform, for running the sequencing and metabolomic analyses. The bioinformatics presented in this paper were carried out using the HPC facilities of the University of Luxembourg (Varrette et al., 2022). In addition, we thank the clinician and hospital personnel that recruited the participants.

This project has received funding from the European Research Council (ERC) under the European Union’s Horizon 2020 research and innovation program (grant agreement No. 863664), and was further supported by the Luxembourg National Research Fund (FNR) CORE/16/BM/11333923 (MiBiPa), CORE/15/BM/10404093 (microCancer/MUST), as well as an “Espoîr en tête” grant from the Rotary Club Luxembourg to P.W. This work was also supported by a Fulbright Research Scholarship from the Commission for Educational Exchange between the United States, Belgium and Luxembourg to P.W. Additional funding was provided by the FNR under INTERMOBILITY/23/17856242.

## Rights Retention Statement

This research was funded in whole, or in part, by the Luxembourg National Research Fund (FNR), grant references CORE/16/BM/11333923 (MiBiPa), CORE/15/BM/10404093 (microCancer/MUST), and INTERMOBILITY/23/17856242. For the purpose of open access, and in fulfilment of the obligations arising from the grant agreement, the author has applied a Creative Commons Attribution 4.0 International (CC BY 4.0) license to any Author Accepted Manuscript version arising from this submission.

## Contributions

Conceptualization P.M, P.W. Patient recruitment, clinical coordination, and sampling: B.M. Bioinformatics, statistics and data visualization: R.V, P.N, P.M. Initial manuscript draft: R.V, P.N. Review and editing: C.C.L., P.M. and P.W. Funding acquisition: C.C.L, P.W. All authors read and approved of the submitted version.

## Notes

### Competing Interest Statement

The authors have declared no competing interest.

