## Supplementary figures and images for "Human gut microbiome gene co-expression network reveals a loss in taxonomic and functional diversity in Parkinson’s disease"

### Supplemental fig.1

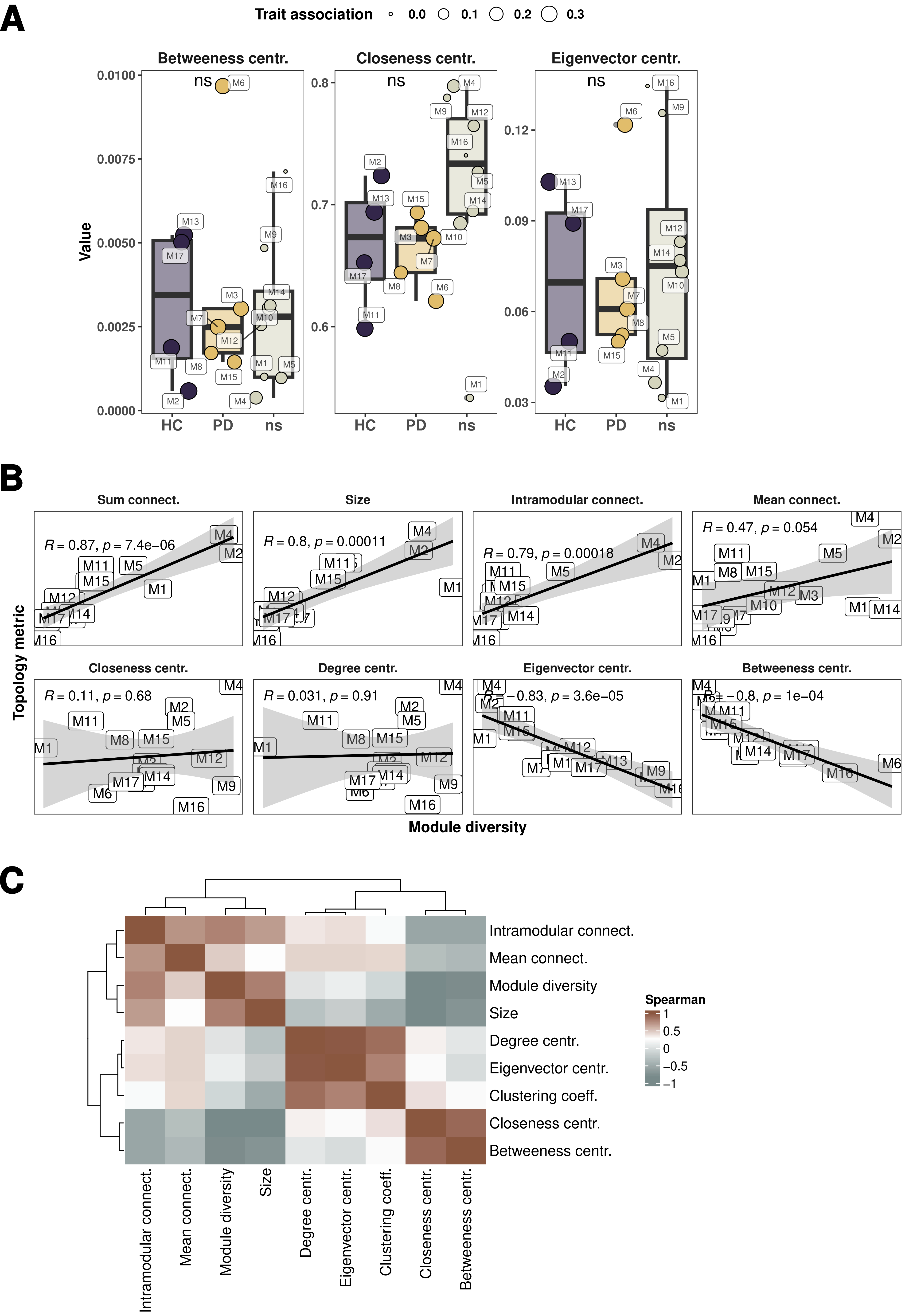

### Supplemental fig.2

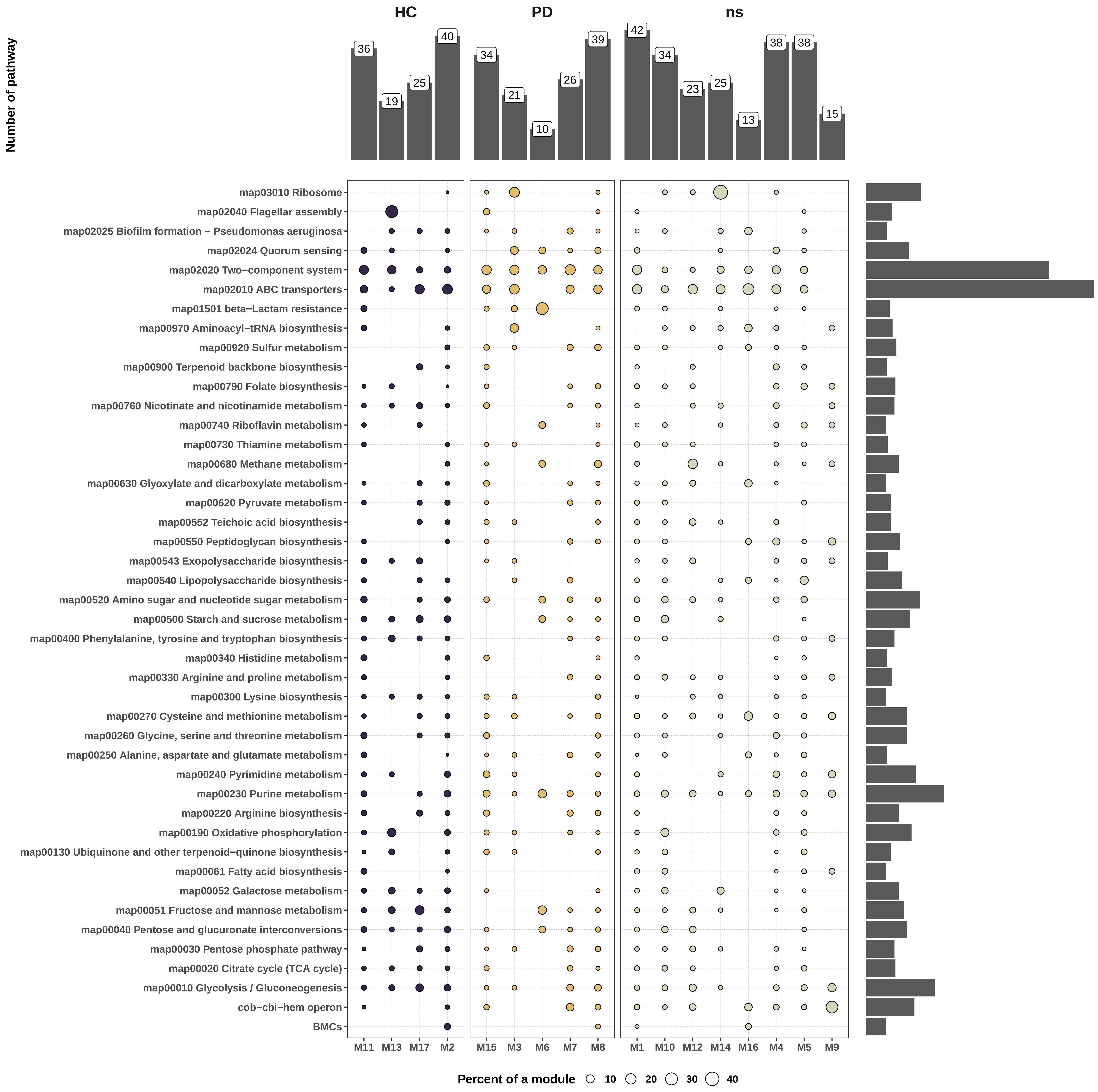

### Supplemental fig.3

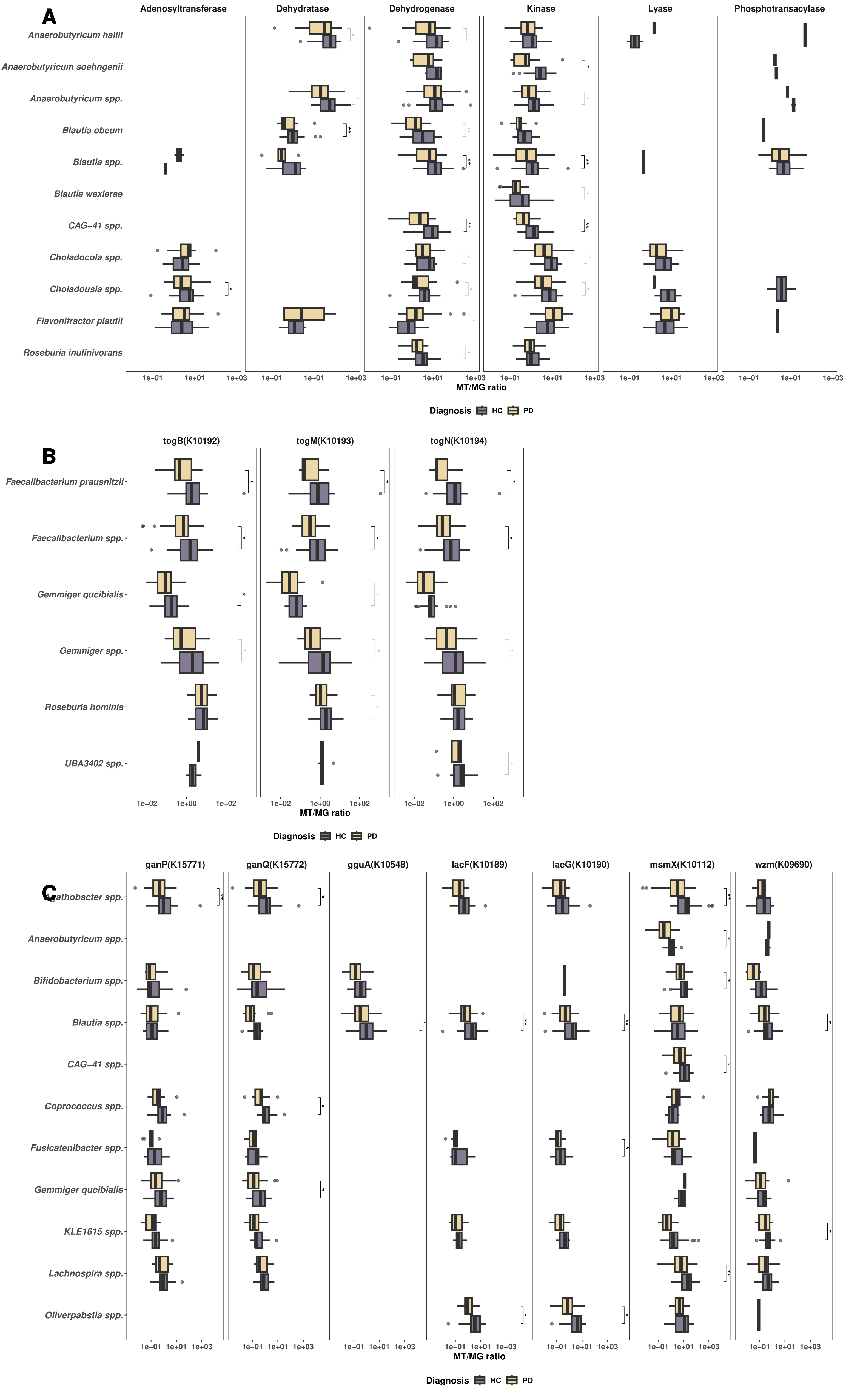

### Supplemental fig.4

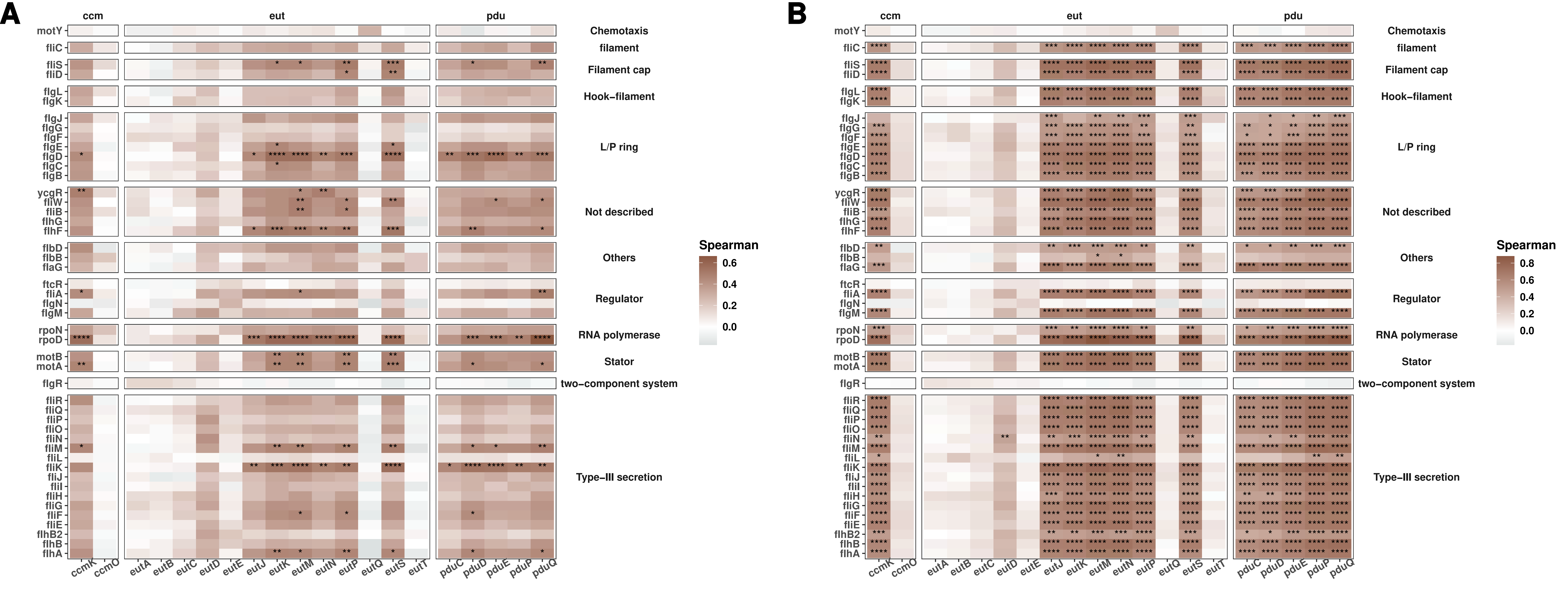
